# *Toxoplasma gondii* infections are associated with boldness towards lions in wild hyena hosts

**DOI:** 10.1101/2020.08.26.268805

**Authors:** Eben Gering, Zachary M. Laubach, Patricia Weber, Gisela Soboll Hussey, Kenna D. S. Lehmann, Tracy M. Montgomery, Julie W. Turner, Wei Perng, Malit O. Pioon, Kay E. Holekamp, Thomas Getty

## Abstract

*Toxoplasma gondii* is widely reported to manipulate the behavior of its non-definitive hosts in ways that promote lethal interactions with the parasite’s definitive feline hosts. Nonetheless, there is a lack of data on the association between *T. gondii* infection and costly behavioral interactions with felids in nature. Here, we report that three decades of field observations reveal *T. gondii* infected hyena cubs approach lions more closely than uninfected peers and have higher rates of lion mortality. Our findings support the hypothesis that *T. gondii’s* manipulation of host boldness is an extended phenotype that promotes parasite transmission from intermediate hosts to feline predators. While upregulating hyena boldness toward lions might achieve this, it may also reflect a collateral influence of manipulative traits that evolved in other hosts (e.g., rodents). In either case, our findings corroborate the potential impacts of a globally distributed and generalist parasite (*T. gondii*) on fitness-related interaction with felids in a wild host.

**One Sentence Summary:** Wild hyenas infected with the parasite *T. gondii* show evidence of costly behavioral manipulation when interacting with lions.

## Introduction

*Toxoplasma gondii* provides an infamous example of putative host-manipulation by a parasite (*1*): within this protist’s diverse warm-blooded hosts, infections are linked to reduced avoidance of, or even attraction to, the odor of feline urine (*2*–*4*). “Fatal attraction” to indirect cues of feline presence is thought to have evolved by natural selection on the parasite to increase trophic (prey to predator) transmission. This could benefit *T. gondii* since the parasite undergoes sexual reproduction within definitive feline hosts to produce recombinant, environmentally stable propagules called oocysts (*5, 6*). Impressively, *T. gondii* also induces other potentially manipulative behaviors in intermediate hosts, including behavioral boldness (*7*). This could similarly promote trophic transmission at the expense of intermediate hosts’ fitness in nature (*8*).

While *T. gondii* is among the best-studied host manipulators, and also causes substantial disease burden in human hosts (*5, 9*), its effects on host behavior have chiefly been studied in laboratory and/or human hosts. A smaller body of research from nature, where *T. gondii* co-evolves with intermediate and definitive hosts, suggests that infection-related behavior might indeed decrease host fitness (*10, 11*). For example, wild-caught rodents harboring naturally-occurring infections exhibit reduced avoidance of odor cues emitted by local felids (*10*), as well as elevated activity levels (*13*), reduced neophobia (*14*), and higher rates of capture in human traps in captive and semi-captive settings (*14*). In wild sea otters, infections are also associated with both neuropathy and shark predation (*15*). However, these studies have not yet demonstrated an association between *T. gondii* infections and naturally occurring behavioral interactions with the parasite’s definitive felid hosts (i.e. vs. experimentally introduced, indirect cues of feline presence). The chief aims of the present study were to test for this relationship, and also examine its potential fitness significance, in a free-living *T. gondii* host.

We used blood samples and detailed field observations spanning three decades to accomplish three goals: 1.) identify demographic, social, and ecological determinants of *T. gondii* infection in a long-lived and highly social host, the spotted hyena (*Crocuta crocuta*); 2.) test whether *T. gondii* infected hyenas exhibit greater behavioral boldness in the presence of lions, and 3.) test whether *T. gondii* infected hyenas have higher rates of lion-inflicted mortality. Our data were collected in a natural setting in Kenya where hyenas frequently interact with lions (*Panthera leo*), which are not only definitive *T. gondii* hosts (*9*) but are also a major source of hyena injuries and mortality. In fact, lions are the leading natural cause of mortality in many hyena populations, including our focal population (*16, 17*). Despite the clear risk lions pose, hyenas engage with them to defend territories, protect relatives, and/or compete for food. Tension between the benefits and costs of these interactions likely explain findings of stabilizing selection on hyena boldness toward lions, favoring individuals with intermediate phenotypes (*18*). This study system therefore permits us to characterize relationships among *T. gondii* infection and naturally occurring behaviors that have well-established fitness consequences for wild hyenas, including behavioral boldness during interactions with lions.

## Results

First, we assessed demographic, social, and ecological determinants of *T. gondii* infection in spotted hyenas. One hundred and nine (109) of 166 surveyed hyenas (65.5%) tested positive for IgG antibodies to *T. gondii*, indicating prior exposure to the parasite. Thirty-six individuals (22%) tested negative, and 21 hyenas (12.5%) yielded results within the “doubtful/uncertain” range of the assay (**Fig. S1**). In keeping with prior studies (*19*), we combined individuals with negative and uncertain diagnoses into a single category, treating them as uninfected in the analysis. A subset of 60 plasma samples were also tested using IFAT diagnostics to confirm consistency between ELISA and IFAT (**Fig. S2;** Pearson’s r(58)=0.70, p<0.001).

Table 1 shows bivariate associations between a) demographic, social and ecological variables (sex, age, dominance rank, and livestock density) and b) prevalence of *T. gondii* infections. We observed no differences in infection prevalence between male vs. female hyenas (61% vs. 69%; *P*=0.33). Hyena cubs (35% infected) had lower infection prevalence than subadults (74%) and adults (80%; overall *P*-difference <0.001). Dominance rank was not associated with the probability of being infected (*P*=0.95). While hyenas sampled in areas of high livestock density showed a trend of higher infection prevalence (79% vs. 62%, *P*=0.08), this association was driven by the overrepresentation of cubs residing in areas with low livestock density, and was not significant after restricting analyses to subadult and adult individuals (*P=0*.*98)*.

Table 1 also summarizes the associations between hyenas’ infection statuses and demographic, social, and ecological variables after adjusting for potential confounding variables. These results recapitulate the outcomes of bivariate tests: neither hyena sex nor dominance rank were associated with infection status (odds ratio [OR] for male vs. female hyenas: 1.11 [95% CI: 0.50, 2.47] and OR for standardized dominance rank 1.11 [95% CI: 0.47, 2.65]). *T. gondii* was more prevalent in older individuals (OR for sub-adults vs. cubs: 5.50 [95% CI: 1.93, 16.92]; OR for adults vs. cubs: 7.92 [95% CI: 3.51, 18.85]), and no difference in prevalence was observed between high vs. low livestock density habitats (OR: 0.47 [95% CI: 0.16, 1.23]).

Second, we investigated associations of *T. gondii* infection with boldness toward lions, as indicated by minimum approach distance to lion(s). **Table S1** shows bivariate associations between a) hyenas’ minimum approach distances toward lions, and b) candidate predictor variables. At alpha=0.05, shorter minimum approach distances were seen in female and older hyenas (i.e., sub-adult or adult), among higher dominance rank hyenas and in areas with high livestock density.

**Table 1.** Prevalence of *T. gondii* infection among 166 spotted hyenas from the Masai Mara, Kenya, and its relationship to demographic and ecological variables.

|  | % (N) |  | OR (95% CI) infected vs. uninfected |  |
| --- | --- | --- | --- | --- |
|  | Uninfected<br>n = 57 | Infected<br>n = 109 | Unadjusted <sup>a</sup> | Adjusted <sup>b</sup> |
| **Sex** |  |  |  |  |
| Female | 31% (30) | 69% (66) | 1.00 (Reference) | 1.00 (Reference) |
| Male | 39% (27) | 61% (43) | 0.72 (0.38, 1.38) | 1.11 (0.50, 2.47) |
| **Age at serodiagnosis <sup>c</sup>** |  |  |  |  |
| Cub (<12 mos) | 65% (32) | 35% (17) | 1.00 (Reference) | 1.00 (Reference) |
| Subadults (12-24 mos) | 26% (9) | 74% (26) | 5.44 (2.15, 14.81)** | 5.50 (1.93, 16.92)** |
| Adult (>24 mos) | 20% (16) | 80% (66) | 7.76 (3.55, 17.80)** | 7.92 (3.51, 18.85)** |
| **Dominance Rank <sup>d</sup>** |  |  |  |  |
| Standardized rank (-1 : 1) | 42% (40) | 58% (56) | 1.02 (0.53, 1.96) | 1.11 (0.47, 2.65) |
| **Livestock density <sup>e</sup>** |  |  |  |  |
| High | 21% (7) | 79% (26) | 1.00 (Reference) | 1.00 (Reference) |
| Low | 38% (50) | 62% (83) | 0.45 (0.17, 1.06)* | 0.47 (0.16, 1.23) |

<sup>a</sup> From a logistic regression model where the independent variable of interest is each socioecological characteristic, and the outcome is infection (yes vs. no).
<sup>b</sup> Mutual adjustment for all demographic and ecological characteristics. In models where sex or human disturbance was the independent variable of interest, age was included as a continuous variable (months).
<sup>c</sup> Assessed on the date the hyena was diagnosed (i.e. the darting date).
<sup>d</sup> Adult female rank or a cubs maternal rank the year during which the hyena lion interaction was observed. On the standardized rank scale, -1 corresponds with the lowest rank and 1 with the highest rank.
<sup>e</sup> Based on illegal livestock grazing in the park during the year in which a hyena was diagnosed.
\* Denotes statistical significance at alpha = 0.10.
\*\* Denotes statistical significance at alpha = 0.05.

Given the age structure of *T. gondii* prevalence in hyenas, coupled with a strong age effect in which older hyenas approach lions more closely than cubs, we conducted separate analyses of behavioral covariates of infection in cubs vs. older individuals. Among cubs, infected individuals had a 47.93 m (95% CI: 16.64, 79.22) shorter minimum approach distance to lions than their uninfected counterparts after controlling for sex and age in months at the time of interaction (**Fig. 1 A, Table 2**). Among subadult and adults, infection was not related to minimum approach distance (1.40 m [95% CI: −3.35, 6.16]; **Fig. 1 B, Table 2**). We further investigated this relationship by limiting our dataset to hyena-lion interactions recorded after the diagnosis dates for seropositive individuals, and prior to the diagnosis dates of seronegative individuals. Still, we observed no association (3.56 m [95% CI: −7.59, 14.71] between infection status and boldness behaviors.

**Table 2.**
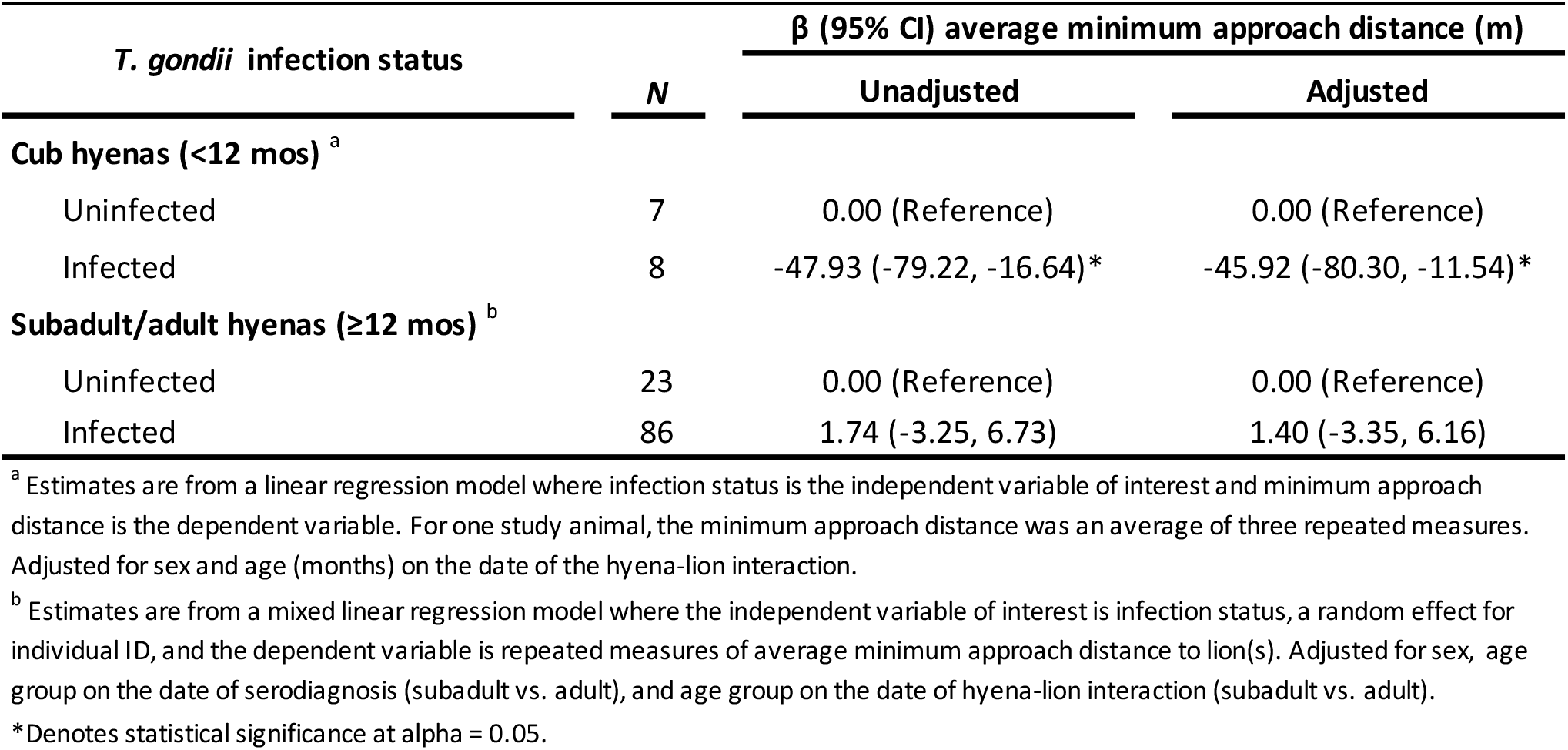
Associations of *T. gondii* infection with minimum approach distance to lion(s) among spotted hyena cubs (N = 15) and subadult/adults (N = 109).

| *T. gondii* infection status | *N* | $\beta$ (95% CI) average minimum approach distance (m) | |
| --- | --- | --- | --- |
|  |  | Unadjusted | Adjusted |
| **Cub hyenas (<12 mos) <sup>a</sup>** |  |  |  |
| Uninfected | 7 | 0.00 (Reference) | 0.00 (Reference) |
| Infected | 8 | -47.93 (-79.22, -16.64)* | -45.92 (-80.30, -11.54)* |
| **Subadult/adult hyenas (≥12 mos) <sup>b</sup>** |  |  |  |
| Uninfected | 23 | 0.00 (Reference) | 0.00 (Reference) |
| Infected | 86 | 1.74 (-3.25, 6.73) | 1.40 (-3.35, 6.16) |

<sup>a</sup> Estimates are from a linear regression model where infection status is the independent variable of interest and minimum approach distance is the dependent variable. For one study animal, the minimum approach distance was an average of three repeated measures. Adjusted for sex and age (months) on the date of the hyena-lion interaction.
<sup>b</sup> Estimates are from a mixed linear regression model where the independent variable of interest is infection status, a random effect for individual ID, and the dependent variable is repeated measures of average minimum approach distance to lion(s). Adjusted for sex, age group on the date of serodiagnosis (subadult vs. adult), and age group on the date of hyena-lion interaction (subadult vs. adult).
\*Denotes statistical significance at alpha = 0.05.

**Fig. 1.**
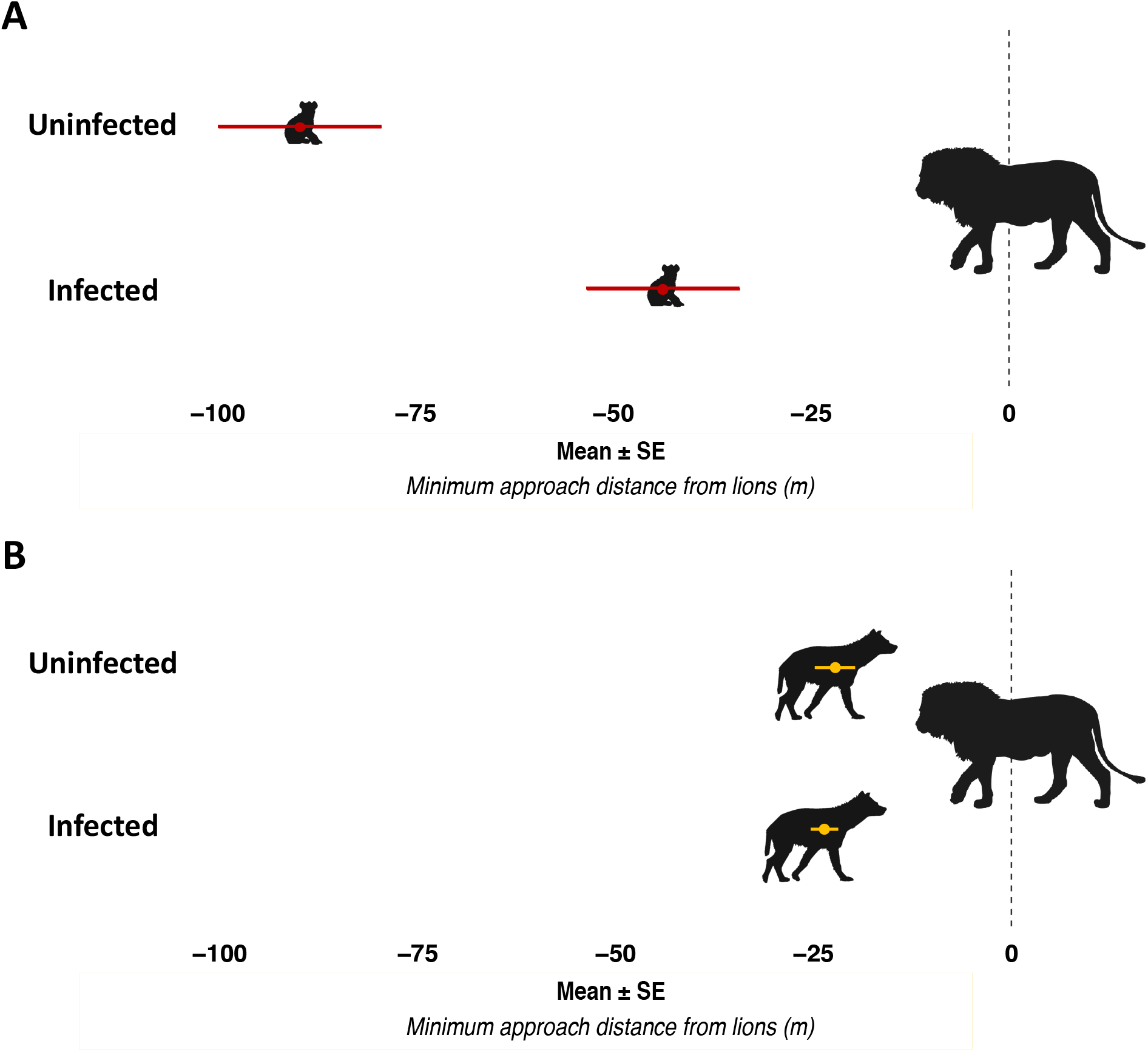
Mean±SE minimum approach distance to lions for seropositive vs. seronegative hyenas based on linear regression models for cubs and mixed-effects linear regression models that included a random intercept for hyena ID to account for correlations among repeated measurements in subadult and adult hyenas. **A**. Among cubs (N=15), models were adjusted for cub sex and age in months on the date of the hyena-lion interaction. **B**. Among subadults/adults (N=109), models were adjusted for hyena sex, hyena age group on the date of diagnosis, and age group on the date of hyena-lion interaction.

In sensitivity analyses, we assessed the potential effects of known determinants of hyena boldness behaviors in this population among cubs, as this was the subgroup within which we found an effect of *T. gondii* on approach distance to lions. First, we included the cubs’ maternal rank as a covariate and noted no appreciable change in the estimate nor the interpretation of results (−29.76 m [95% CI: −55.95, −3.57]). Next, we adjusted for the presence of a male lion during the hyena-lion interaction (−47.71 m [95% CI: −84.10, −11.33]). Finally, we accounted for the presence of food during the interaction (−37.24 m [95% CI: −66.62, −7.87]). None of these adjustments markedly changed the direction, magnitude, or precision of the estimate for *T. gondii* infection status in relation to approach distance.

Third, we explored associations of *T. gondii* infection with lion-related mortality. Among 33 mixed-age hyenas with known mortality causes, infected hyenas were nearly twice as likely to die by lions than by other known causes (52% vs. 25%). When we assessed this in regression models, infected hyenas were 3.85 (95% CI: 0.68, 32.46; *P*=0.16) times more likely to die by lions than uninfected animals after accounting for sex, though this effect was not significant (**Table 3**). Among hyenas infected as cubs, 100% of the deaths were caused by lions, while only 17% of deaths of uninfected cubs were caused by lions. While the cubs diagnosed as part of this analysis were not all from the same litter or birth cohort, they were sampled during a single early period of our study from 1990-1999. Thus, in the small subsample of 11 cubs, the probability of dying by lions vs. other known sources of mortality was greater among infected than uninfected individuals (Fisher’s Exact Test *P*=0.01).

**Table 3.**
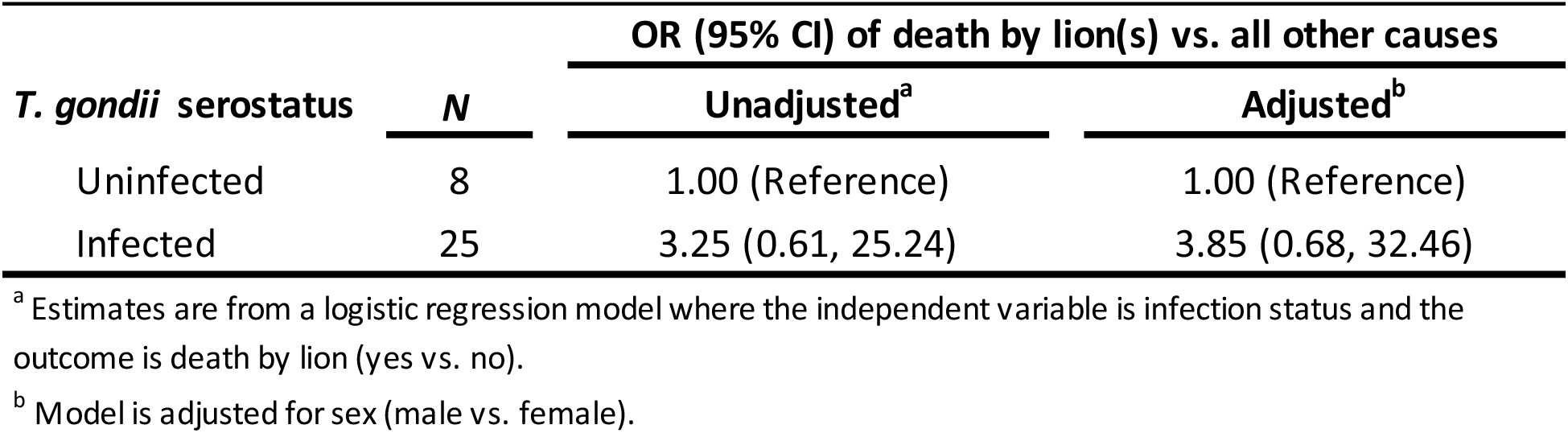
Associations of *T. gondii* infection with odds of death by lion(s) among 33 hyenas.

| *T. gondii* serostatus | *N* | OR (95% CI) of death by lion(s) vs. all other causes |  |
| --- | --- | --- | --- |
|  |  | Unadjusted<sup>a</sup> | Adjusted<sup>b</sup> |
| Uninfected | 8 | 1.00 (Reference) | 1.00 (Reference) |
| Infected | 25 | 3.25 (0.61, 25.24) | 3.85 (0.68, 32.46) |

<sup>a</sup> Estimates are from a logistic regression model where the independent variable is infection status and the outcome is death by lion (yes vs. no).
<sup>b</sup> Model is adjusted for sex (male vs. female).

## Discussion

In this study, *T. gondii* infection was associated with behavioral boldness that brought infected hyena cubs into closer proximity to lions, thereby increasing risk of mortality by lions. The fact that we saw no association between the infection status of older (subadult and adult) hyenas and approach distance to lions may indicate that experienced individuals better assess threats and inhibit risky behavior. Testing this, and other potential causes of age-dependency in infection-related behavior will require additional data. We also noted that infected hyenas were more likely to be killed by lions than by other causes, although this effect was most pronounced in cubs. Together, our results provide evidence for a link between *T. gondii* infection, boldness toward lions, and fitness in a wild population of hyenas. This link is mechanistically plausible given that lions readily attack and kill hyenas (*17, 20*) and are a leading source of hyena mortality in the wild (*16*). Hyenas with above-average boldness in the presence of lions also have reduced longevity, which may result from injuries and lethal wounds inflicted by lions (*18*).

Our results also suggest that the prevalence of *T. gondii* infection is influenced by demographic factors. For example, older hyenas were more likely to be infected. A recent survey of carnivores from the Serengeti ecosystem found similar age-structured prevalence of *T. gondii* infection in spotted hyenas, and suggested that ingestion of infected prey or carrion – of which older individuals have a longer cumulative exposure over the life span – may be an important source of infection (*21*).

Against our predictions, hyenas living where there are greater amounts of illegal grazing of domestic livestock within the Masai Mara Reserve do not exhibit higher prevalence of *T. gondii* infection. Originally, we hypothesized that carnivores face greater risk of infection within close proximity of domesticated animals and human commensals (e.g., feral cats and rodents) that serve as disease reservoirs. Indeed, agricultural activity is a known amplifier of *T. gondii* prevalence in livestock, human commensals, and wild animals (*22, 23*), and freshwater runoff from urbanized areas is associated with higher disease prevalence in sea otters (Miller et al. 2002). We had therefore predicted that the more recent samples from localities undergoing agricultural intensification would exhibit higher *T. gondii* prevalence. However, we found that *T. gondii* was equally prevalent in hyenas sampled decades ago as in hyenas sampled more recently, and also equally prevalent in regions of the Masai Mara that are either inundated with or protected from livestock grazing. This indicates the potential for historical co-evolution between *T. gondii* and its hosts within this system, and also the parasite’s widespread distribution throughout the Mara.

It is tempting to speculate that our findings reflect an underlying mechanism through which manipulating hyena boldness promotes *T. gondii* transmission to lions. As a viable definitive host, lions are capable of shedding large numbers of recombinant, environmentally stable *T. gondii* oocysts into the local environment. These oocysts can infect a multitude of warm-blooded host species that inhabit the Mara landscape, including hyenas. However, it is noteworthy that lions rarely consume hyenas after killing them. Although this reduces trophic transmission opportunities, killings do involve significant exchanges of blood and tissue that could feasibly transmit *T. gondii* (**Movie S1**). Additionally, hyenas killed by lions are typically consumed by a variety of carrion-feeding mammals and birds, including many intermediate *T. gondii* hosts (*24*). Given that a single hyena carcass can infect many highly mobile carrion feeders (e.g., vultures), these scavengers may play important roles in amplifying and dispersing the parasite beyond the capacity of terrestrial hosts.

In an alternative, highly plausible scenario, the behavioral phenotypes of infected hyenas may simply represent “collateral manipulation” via traits that evolved to influence other host species like rodents. This scenario appears to manifest in *T. gondii-*infected humans, who exhibit riskier behavior (*5*) despite being dead-end hosts. This possibility is further supported by findings that homologous neural and hormonal regulators of behavior are similarly altered by *T. gondii* infections of human and non-human hosts (*25*). However, the concept of collateral manipulation has not yet been well-integrated into studies of wild animal populations, where researchers can directly assess its ecological and evolutionary significance.

This study is not without limitations. For instance, hyenas’ behaviors towards lions may not be independent of one another given the social nature of this species. An ideal analytical strategy to reflect this would involve use of hierarchical models to cluster by observation session to account for correlations among hyenas observed together during their interaction with lions. At present, we are underpowered to conduct such an analysis. Furthermore, a more sophisticated measure of potential modes of transmission among livestock, pastoralists and hyenas are required to better understand infection risk as a product of humans and wildlife living in close proximity. This would involve ascertaining incident infection to more precisely estimate when a hyena first contracts the parasite. Future studies with appropriate data are warranted.

In summary, our results suggest the *T. gondii* infection is likely deleterious to hyenas, at least when contracted early in life. If similar effects occur in other *T. gondii* hosts, the ecological and evolutionary significance of this globally abundant, highly generalist protist may be vastly underappreciated. We encourage further explorations of fitness-related behavior in natural settings – both for infections of *T. gondii*, which involve a substantial proportion of earth’s mammals and birds, and for other parasites suspected to alter host behavior to serve their own evolutionary interests.

## Acknowledgements

We thank the Kenyan National Commission for Science, Technology, and Innovation, the Naboisho Conservancy, the Narok County Government, the Kenya Wildlife Service and Brian Heath for permission to conduct research in the Mara ecosystem. We also thank Chiara Bowen and Nichole Grosjean for lab assistance, Samantha Gregg, Katie Keyser, Leah McTigue, and Abigail Thiemkey for assistance extracting distances between lions and hyenas, the NSF-BEACON center for the Study of Evolution in Action for funding, the Mara Hyena Project field crew, and the residents of the Mara ecosystem for their direct and indirect support of long-term field research in their backyard. We thank members of the Getty and Holekamp labs for their feedback on this project and paper.

## Author contributions

Conceptualization, EG, ZML; data generation and curation, EG, ZML, PW, GSH, KDSL, TMM, JWT, MOP; formal analysis, ZML, WP; funding acquisition, EG, ZML, KEH; investigation, EG, ZML; methodology, EG, ZML, PW, GH, KDSL, TMM, WP; resources, EG, PW, GSH, KEH; supervision, KEH, TG; visualization, EG, ZML; writing – original draft, EG, ZML; writing – review & editing, EG, ZML, PW, GSH, KDSL, TMM, JWT, WP, MOP, KEH, TG.

## Data and materials availability

Upon publication, all data and analysis code will be made available through public GitHub pages (https://github.com/laubach and https://github.com/MaraHyenaProject) as well as on DRYAD.

