## Supplemental Material for "*Toxoplasma gondii* infections are associated with boldness towards lions in wild hyena hosts"

**This PDF file includes:**

Materials and Methods
Figs. S1 to S2
Tables S1
Caption for Movies S1

**Other Supplementary Materials for this manuscript include the following:**

Movies S1

### Materials and Methods

The Mara Hyena Project

This study uses data and samples from the Mara Hyena Project, a long-term field study of individually known spotted hyenas that have been observed since May 1979. Study hyenas are monitored daily and behavioral, demographic, and ecological data are systematically collected and entered into a database. Here, we used data from four different hyena groups, called clans, as well as historic information about ecological conditions in the Masai Mara National Reserve. We maintained detailed records on the demographics of our study population, including sex, age, and the dates of key life-history milestones such as birth, weaning, dispersal and death. In the ensuing sections, we describe data collection and data processing procedures for assessment of *T. gondii* infection diagnosis, quantification of demographic and ecological determinants of infection status, and assessment of behavioral (boldness) and fitness (cause of mortality) characteristics hypothesized to be a consequence of positive *T. gondii* infection. The present analysis includes 168 hyenas, but specific subsamples vary depending on the particular hypothesis being tested.

Biospecimen collection and assessment of Toxoplasma gondii exposure

As part of our long-term data collection, we routinely darted study animals in order to collect biological samples and morphological measurements. Of special relevance to this study is our blood collection procedure. We immobilized hyenas using 6.5 mg/kg of tiletamine-zolazepam (Telazol ®) in a pressurized dart fired from a CO2 powered rifle. We then drew blood from the jugular vein into sodium heparin-coated vacuum tubes. After the hyena was secured in a safe place to recover from the anesthesia, we took the samples back to camp where a portion of the collected blood was spun in a centrifuge at 3000 rpm for 10 minutes to separate red blood and white blood cells from plasma. Plasma was aliquoted into multiple cryogenic vials. Immediately, the blood derivatives, including plasma, were flash frozen in liquid nitrogen where they remained until they were transported on dry ice to a -80°C freezer in the U.S. All samples remained frozen until time of laboratory analysis for the *T. gondii* assays.

Using archived plasma, we diagnosed individual hyenas using the multi-species ID Screen® Toxoplasmosis Indirect kit (IDVET, Montpellier). This ELISA-based assay tests for serological (IgG) reactivity to *T. gondii's* P-30 antigen and has been used in many prior studies of *T. gondii* in diverse mammals (*1*). The output of the assay is an SP ratio, which is calculated as colorimetric signal of immunoreactivity for a tested blood sample (S) divided by that of a positive control (P), after subtracting the background signal for the ELISA plate (i.e. a negative control) from both S and P. We tested 168 plasma samples from spotted hyenas and determined infection status based on the kit manufacturer's criteria for interpreting S/P: <40% = negative result, 40%-50% = doubtful result, >50% = positive result. Only 21 hyenas (13%) fell within the "doubtful" range of ELISA S/P values. We treated these individuals as negative in our analyses, following protocols from other recent studies (*1*). Although ELISA-based assays performed relatively well in prior studies of *T. gondii* in both hyenas (*2*) and other mammals (*1*), the method can also be sensitive to cross-reactivity with antibodies to other, related parasites (*3*). We therefore retested 60 randomly chosen plasma samples using an alternative, indirect fluorescence agglutination test (IFAT), which is less sensitive to cross reactivity (*2*), in order to confirm that the two methods yielded similar results using our plasma samples. Samples were submitted to the Michigan State University Veterinary Diagnostic Laboratory for a standard diagnostic IFAT procedure that used whole organism, from reagents supplied by VMRD, Pullman, WA. In brief, the IFAT measures the maximum dilution of a plasma sample at which immunoreactivity to *T. gondii* antigen is visible by microscopy. Two samples were excluded from our main analyses because of suspected assay error (e.g. one negative SP ratio and one additional IFAT vs. SP ratio discrepancy) making our final diagnostic sample size, N=166 hyenas. However, it should be noted that inclusion of these two questionable data points did not substantively change our results and had no effect on our analyses of hyena boldness or fitness as these two hyenas lacked data required for those analyses. Diagnostic assays were performed by people who were blind to the individual hyena's demographic, ecological, and behavioral data.

Demographic, social, and ecological characteristics

The first aim of this analysis was to identify demographic and ecological correlates and determinants of *T. gondii* infection. The key characteristics of interest include sex and age, two demographic traits that have previously been implicated in health and behavioral outcomes, as well as exposure to livestock density.

Sex. We determined the sex of each hyena based on the glans morphology of its erect phallus during field observations; this is reliable starting at 3 months of age (Frank, Glickman, & Powch, 1990).

Age. We aged each hyena by back-calculating its birthdate based on its physical appearance when first observed in infancy. Based on its pelage, morphology and behavior, we are able to determine a birthdate age with an accuracy of ± 7 days (*5*). We used this method to determine each hyena's age in months at the time of blood collection. In the analysis, we assessed age continuously in months, as well as in distinct age groups divided into cubs (<12 months), subadults (>12 to ≤24 months), and adults (>24 months of age). The age cut-offs were determined based on the timing of major life history milestones; weaning occurs at approximately 12 months of age and hyenas of both sexes achieve reproductive competence at around 24 months of age (*5*, *6*).

*Dominance Rank.* As part of our routine data collection, all aggressive interactions between hyenas are recorded, such that we can calculate rates of threat displays, chases, and bites between clan members. Agonistic interaction data are used to calculate each hyena's dominance rank each year via a matrix of dyadic wins and losses in fights (*7–9*). Values from each rank matrix are normalized as a continuous variable from -1 (lowest) to 1 (highest) and are regularly updated to account for demographic change. We use maternal rank as a proxy of each cubs rank until they learn their own rank.

*Livestock density*. To quantify exposure to livestock density, we took advantage of naturally occurring variation in exposure to human activity across distinct regions of the Reserve. We classified all hyenas from the western region of the reserve (also known as the Mara Triangle) as "low livestock density" due to strict bans on livestock grazing and travel on foot in this area. The eastern side of the Reserve borders pastoralist villages that have experienced an increase in human population growth in recent years, especially around the burgeoning Talek community (*10*). Additionally, assessment of trends in livestock counts within the reserve indicates a marked increase in illegal livestock grazing on the eastern side of the reserve starting in 2000, followed by another increase between 2009 and 2013. These changes coincided with parallel shifts in hyena demography and wildlife community composition (*11*). To improve discriminatory power in our analyses that reflect these changes in livestock density and shifts in ecology, we enriched our sample selection to include hyenas from areas of low and higher livestock density. For "low livestock density," we selected animals from the eastern side of the reserve sampled before 2000 along with animals from the western side of the reserve (any time period). For "high livestock density," we selected animals from the eastern side of the reserve from 2012 onward.

Boldness behaviors and fitness

In addition to identifying determinants and correlates of *T. gondii* infection, we also sought to explore the effects of infection status on hyena behavior and fitness. Over the duration of our study, we documented all observed hyena-lion encounters i.e., all instances where at least one hyena and at least one lion approached to within approximately 200m of one another. In 731 observation sessions we recorded 3791 minimum distance estimates between individual hyenas and one or more lions along with the date, location, and identities of all hyenas present, as well as whether food (a dead prey animal or its components) or a male lion was present during the encounter, as both these factors are known to influence hyena behaviors. All boldness behaviors were extracted by four individuals blinded to infection status with 83% agreement across seven metrics recorded during hyena-lion interactions(*12*).

*Minimum approach distance to lion(s)*. Based on previous findings in this study population that minimum distance from lions is a measure of boldness that shows inter-individual consistency and corresponds with fitness(*13*), we used this metric as an indicator for behavioral boldness. During each lion-hyena encounter, we recorded the distances between lions and individual hyenas in meters using 20-minute scan sampling of individual hyena distances from the nearest lion, as well as all-occurrence sampling of close behavioral interactions between lions and hyenas (e.g., a hyena comes within 10m of a lion). Because the body length of an adult hyena is approximately 1m, we are able to accurately estimate approach distances at this scale.

Due to the inherent frenetic activity at some hyena-lion encounters, some of the minimum approach distances were recorded as ranges (e.g. 10-15m) or inequalities (e.g., <10m). For ranges, we calculated the mid-point and used this value in the analysis (e.g. if the range was 10-15m, then we used 12.5m in our calculations). If the distance range was large (>25m) and therefore highly uncertain (i.e. if the range exceeded ½ a standard deviation as estimated from the minimum approach distance data set), we removed it from the dataset. Of the 529 approach distance ranges in our data set, 225 were removed because the range exceeded 25m. We retained inequality distances by using the 'less than' distance if the recorded distance was smaller than 25m (approximately the mean [mean=45m] minus ½ a standard deviation [sd=50m] of all hyena minimum approach distances) and by including the 'greater than' distances if the recorded distance was greater than 75m (approximately the mean plus ½ a standard deviation of hyena minimum approach distances). For example, a distance recorded as <50 m would be removed from the data set as it could include a wide range of actual distances (0-50 m), while a recorded distance of <15 m was retained in the data set as 15 m. As a result of filtering inequality distances with large uncertainty, we removed 67 of 72 approach distances recorded as inequalities. Finally, we filtered the hyena approach distance to lions by removing instances when the minimum approach distance exceeded 100m, given that at this range hyenas and lions pose little threat to one another. After filtering, our final data set included 2,724 minimum approach distance estimates. It should be noted that during any particular hyena-lion interaction, we retained a single minimum approach distance for each hyena, but over their lifetime hyenas interact with lions on multiple occasions, thus the repeated minimum distance measures for hyenas.

*Death by lion(s)*. Our longstanding behavioral database also documents the source of mortality for each hyena whenever known. Deaths attributed to lions included cases in which lions were observed killing hyenas and when fresh corpses of hyenas were found with puncture wounds in each corpse made by canine teeth that were too far apart to have been inflicted by anything but a lion. In our analysis, we dichotomized cause of mortality as death by lion vs. all other causes and evaluated this as a binary outcome in the statistical analysis.

Statistical analyses

In the analysis, we tested three hypotheses: (***H1***) higher livestock density is associated with higher risk of *T. gondii* infection in spotted hyenas; (***H2***) infected hyenas behave more boldly towards lions than uninfected hyenas, as indicated by a shorter minimum approach distance; (***H3***) *T. gondii* infection imposes fitness costs on the host, as indicated by greater odds of death by lion(s). We describe methods for testing each hypothesis below, following a description of our general approach to data analysis.

*General approach*. Prior to formal analyses, we assessed the distributions of all variables. This included viewing the distributions and calculating descriptive statistics for continuous variables (e.g., minimum approach distance towards lions) to check for deviations from normality and missing values. We also assessed frequency distributions for all categorical variables (infection status, sex, age group, food presence, and livestock density). Finally, as part of our data exploration, we conducted bivariate analyses of associations between demographic and ecological characteristics and infection status, as well as associations of demographic or ecological characteristics with behavioral outcomes to identify covariates for inclusion in multiple variable analysis. In all models, we considered an estimate to be statistically significant at a nominal cut-off of alpha=0.05, but also noted those that were significant at alpha=0.10 given our relatively small sample sizes. Data cleaning and analyses were performed in program R version 3.6.2. Linear mixed models were conducted using the lme4 package, version 1.1.21 and least squared means estimates were calculated using SAS (version 9.4; Cary, NC, USA)

*H1: Greater livestock density is associated with higher risk of T. gondii infection.* In this portion of the analysis, we used univariable logistic regression to investigate the relationship between livestock density (high vs. low) as the primary explanatory variable of interest, and *T. gondii* infection (positive vs. negative) as the dependent variable. In addition, we also explored associations of other key demographic characteristics as determinants of infection, namely sex and age at diagnosis (in months). Following the univariable analysis, we also examined mutually-adjusted associations among the above variables. In models where sex and livestock density were the independent variables of interest, age was entered as a continuous variable (months) to improve statistical power.

*H2: Infected hyenas behave more boldly towards lions, as indicated by shorter minimum approach distances.* To investigate the extent to which infection status is related to boldness behaviors, we used simple and multiple-variable linear regression models in which *T. gondii* diagnosis (infected vs. uninfected) was the independent variable of interest, and hyenas' average minimum approach distance (m) was the outcome. We stratified all models by age group such that cubs were analyzed separately from subadults and adults. We made this decision based on bivariate associations that revealed a significant age structuring of infection status (i.e., much higher prevalence of infection in subadults and adults than in cubs), as well as significant effects of age on hyena approach distances towards lions (i.e., older hyenas were much more likely to approach lions closer than younger hyenas). The cub models included individual hyenas that had both infection diagnosis and hyena-lion interaction data during their first year of life. Similarly, the subadult and adult models were restricted to include only infection diagnosis and hyena-lion interactions collected from hyenas 12 months of age and older.

When exploring associations among cubs, we first examined the unadjusted association of *T. gondii* diagnosis with minimum approach distance to lions. Next, we controlled for sex, age in months on the date of the interaction with lions, and whether or not food was present during the interaction. The age distributions of hyena cubs during observed interactions with lions were 2.7-8.5 months for uninfected cubs and 3.2-11.8 months for infected cubs. We did not need to account for livestock density because all cubs were sampled in low livestock density areas. Additionally, for all but one cub, we only had a single minimum approach distance recorded to lions. For the cub with multiple measures (N=3), we took the average of its minimum approach distances for use in the analysis.

When exploring associations among subadults and adults, we used a similar modeling strategy to that used for cubs, except rather than using conventional linear regression, we employed mixed linear regression models to account for the multiple assessments of minimum approach distance to lions for hyenas in these two age groups (median=5 measurements per hyena) via a random intercept for the hyena's study ID. After examining unadjusted associations, we implemented a multiple variable model that adjusted for age group (subadult vs. adult), sex (male vs. female), food present during the interaction (yes vs. no), and livestock density during the year of the interaction (high vs. low).

In addition to the above analysis for subadult and adult hyenas, which leveraged all available hyena-lion interaction data, we also conducted sensitivity analyses on a restricted data set wherein we only considered approach distances from lions that occurred prior to the diagnostic date among hyenas who tested negative for *T. gondii* infection, thus ensuring these represented behaviors of uninfected hyenas. Similarly, we only considered hyena-lion interaction data that occurred after the diagnostic date for individuals who tested positive for *T. gondii* infection. The rationale for this approach is rooted in achieving temporal separation to avoid erroneously examining hyena-lion interactions for negative diagnosis hyenas who subsequently became infected and vice versa (*nota bene*: we did not do this for cubs given our small sample size in this age group and because the small age range limited the possibility that a hyena's approach from lions did not reflect its infection diagnosis). Using this restricted dataset, we modelled the associations between infection status and each hyena's closest approach distance to lions following the previously described modeling strategy.

*H3: T. gondii infection imposes fitness costs on the host, as indicated by greater odds of death by lion(s)*. Here, we assessed the probability that *T. gondii* infection in hyenas was associated with lion-induced mortality. To do this, we used logistic regression models to compare the odds of mortality due to lions vs. all other known causes of mortality for infected vs. uninfected hyenas. Following unadjusted analysis, we controlled for sex in a multiple-variable logistic regression analysis. Due to small sample sizes (i.e. cells in cross tabulations with N=0) we were not able to adjust for hyenas' ages and livestock density levels. However, we were able to use a two-by-two table and Fisher's exact test to determine whether the probability of dying by lions vs. other sources of mortality differed between infected and uninfected cubs.

*Sensitivity analyses*. Dominance rank, presence of food, and presence of male lions are key determinants of boldness behaviors in hyenas. Therefore, in age-stratified subgroup analyses where *T. gondii* infection was a significant determinant of approach distance to lions (i.e., in cubs only), we included maternal rank, presence of food, and presence of male lions to rule out the possibility of extraneous causes of boldness behavior. We then assessed the extent to which inclusion of each of these variables, singly, changed the direction, magnitude, and precision of the estimate for *T. gondii* infection in relation to approach distance to lions.

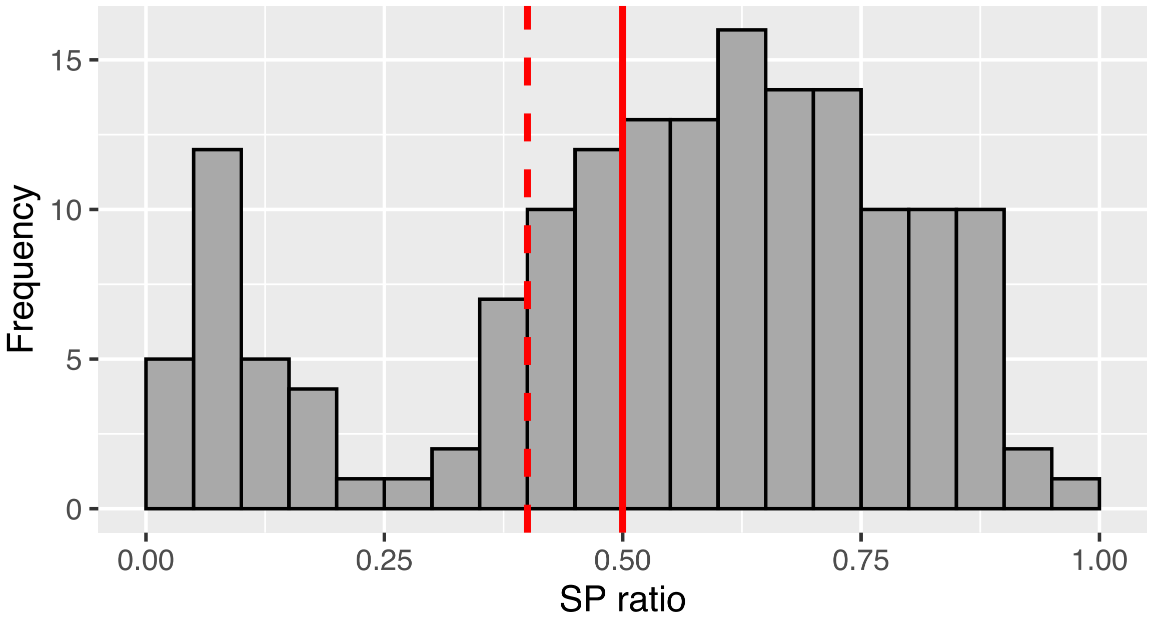


**Fig. S1.** Distribution of SP ratios (from ELISA assays of *T. gondii* antibodies) from spotted hyenas sampled in or near Kenya's Masai Mara National Reserve. The dashed red line is the upper SP ratio cutoff for negative diagnoses, and the solid red line is the lower cutoff for positive diagnosis. The region between the red lines corresponds to the SP ratio range for uncertain, which were treated as negative in this study.

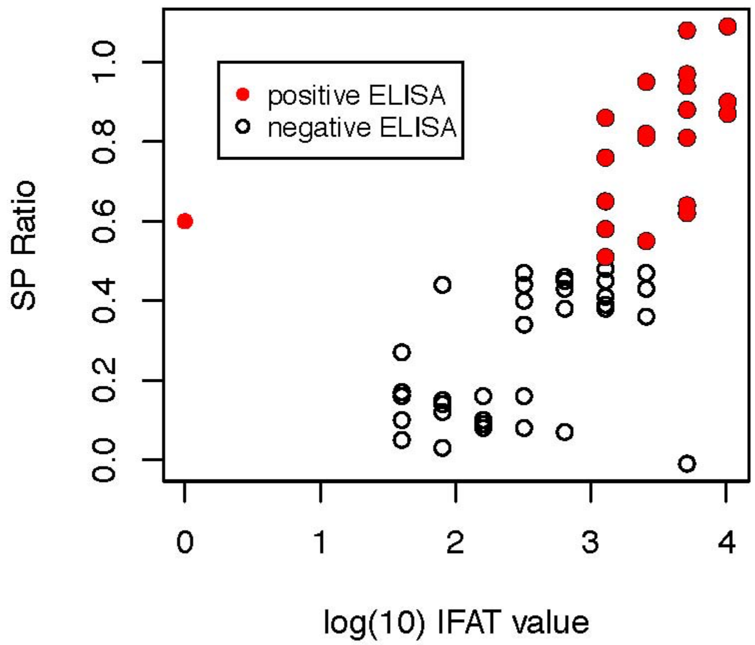


**Fig. S2**. Correspondence between ELISA and IFAT scores for plasma samples from 60 individual hyenas, which were tested using both methods to help corroborate our ELISA diagnostic methods.

**Table S1.** Mean ± SE of average minimum approach distance to lion(s) within categories of demographic and ecological characteristics among 156 hyenas from the Masai Mara, Kenya.

| | Mean ± SE[a] minimum approach distance to lion(s) (m) | *P*-value[b] |
|---|---|---|
| **Sex** | | 0.049 |
| Female | 24.20 ± 1.08 | |
| Male | 27.94 ± 1.57 | |
| **Age at the time of hyena-lion interaction** | | <0.0001 |
| Cub (<12 mos) | 49.62 ± 4.98 | |
| Subadults (12-24 mos) | 23.27 ± 2.40 | |
| Adult (>24 mos) | 24.92 ± 0.96 | |
| **Dominance Rank** [c] | | |
| Per 1 unit increment in standardized rank | -5.14 ± 1.76 | 0.037 |
| **Livestock density** [d] | | 0.003 |
| High | 19.79 ± 2.10 | |
| Low | 26.86 ± 1.11 | |

[a] Estimates are from a mixed linear regression model where the independent variables include the characteristic of interest and a random intercept for hyena ID, and the dependent variable is repeated assessments of a hyena's minimum approach distance from lion(s).

[b] From an independent t-test for sex, food presence, and human disturbance; from a Wald chi-squared test for age group.

Minimum approach distances reprersent repeated measurements from some hyenas; total number of distance measurements = 1,190.

[c] Adult female rank or a cubs matnernal rank the year during which the hyena lion interact was observed. On the standardized rank scale, -1 corresponds with the lowest rank and 1 with the highest rank.

[d] Based on illegal livestock grazing in the park during the year in which a hyena was diagnosed.

**Movie S1.**

Sequence of a hyena being killed by a lion within the study's focal population (in Kenya's Masai Mara region). Although lions rarely consume hyenas after killing them, the video demonstrates a potential for *T. gondii* transmission via the ingestion of infectious blood and tissue. Footage collected by Malit Pioon April 7, 2019.
